# Moana: A robust and scalable cell type classification framework for single-cell RNA-Seq data

**DOI:** 10.1101/456129

**Authors:** Florian Wagner, Itai Yanai

## Abstract

Single-cell RNA-Seq (scRNA-Seq) enables the systematic molecular characterization of heterogeneous tissues at an unprecedented resolution and scale. However, it is currently unclear how to establish formal cell type definitions, which impedes the systematic analysis of scRNA-Seq data across experiments and studies. To address this challenge, we have developed *Moana*, a hierarchical machine learning framework that enables the construction of robust cell type classifiers from heterogeneous scRNA-Seq datasets. To demonstrate *Moana*’s capabilities, we construct cell type classifiers for human immune cells that accurately distinguish between closely related cell types in the presence of experimental perturbations and systematic differences between scRNA-Seq protocols. We show that *Moana* is generally applicable and scales to datasets with more than ten thousand cells, thus enabling the construction of tissue-specific cell type atlases that can be directly applied to analyze new scRNASeq datasets. A Python implementation of *Moana* can be found at https://github.com/yanailab/moana.

## Introduction

Single-cell RNA-Seq is a breakthrough technology with applications in diverse areas of biological research^1–3^. Thanks to exponential increases in throughput over the last years, current protocols enable the efficient processing of more than ten thousand cells in a single experiment^4^. Many studies have demonstrated the unprecedented power of scRNA-Seq to provide systematic views (sometimes referred to as “atlases”) of cell type heterogeneity across organisms, tissues, and developmental stages, which in many cases has led to reports of novel cell types or states^5–8^.

Despite these advances, it remains unclear how to formally define cell types based on scRNA-Seq data, a problem that is complicated by high levels of technical noise^9–12^. In the absence of formal cell type definitions, the analysis of each new scRNA-Seq dataset requires the manual assignment of cell type identities. This laborious task is affected by a large number of study-specific factors, including the choice of clustering method and associated parameters^13^, the technical quality of the data, the set of potential marker genes considered for each cell type, etc. Consequently, different scRNA-Seq studies of the same tissue may rely on inconsistent cell type definitions, which makes it difficult to compare and synthesize study results. Similarly, it is difficult to conduct scRNA-Seq follow-up studies of specific cell types without an unbiased method to re-identify these cell types in a new dataset.

The definition of cell types can be properly framed in machine learning terms as a *classification* problem. In other words, if there existed an algorithm that could accurately and reliably predict (assign) cell type identities to cells in a new scRNA-Seq dataset, then this algorithm would embody a quantitative, expression-based definition of those cell types. However, a generally applicable machine learning framework for scRNA-Seq cell type classification must address a formidable array of challenges: First, it must overcome the high levels of technical noise inherent to scRNA-Seq data^10–12^. Second, it must be robust to systematic differences between datasets generated with different scRNA-Seq protocols^14^. Third, it must be able to accommodate a potentially large number of cell types and closely related subtypes that are commonly present in complex tissues. Fourth, it must be capable of processing large datasets consisting of tens of thousands of cells in reasonable amounts of time, and using limited computational resources (e.g., memory). Finally, it must provide a systematic approach to assess classification performance in the absence of a ground truth, as scRNA-Seq datasets for experimentally purified subpopulations are frequently unavailable.

To address some of the challenges arising from the lack of formal cell type definitions, methods that facilitate the “alignment” and joint clustering analysis of multiple scRNA-Seq datasets have been developed^14,15^. These approaches were shown to successfully overcome batch effects, and in principle allow the cell type annotations of one dataset to serve as a reference for analyzing other scRNA-Seq datasets from the same tissue. However, as these methods lack explicit representations of cell types, they do not obviate the need to conduct a manual clustering analysis, and thus do not eliminate the risk for study-specific biases to arise. An early scRNA-Seq study applied a multinomial mixture model to study the response of specific mouse immune cell populations following exposure to lipopolysaccharide, but did not quantify classification performance^16^. A recent study proposed an algorithm termed *scmapcluster*^17^ that “projects” cell types across scRNA-Seq datasets by selecting informative genes and assigning cell identities to individual cells based on their correlation with average cell type expression profiles. While this represented an innovative classification approach for classifying scRNA-Seq data, validations were largely restricted to pancreas tissue composed of highly specialized cell types, and the ability to classify cells belonging to more closely related cell types remained unclear.

Here, we describe a new machine learning framework, termed *Moana*, which enables the construction of robust cell type classifiers from heterogeneous scRNA-Seq datasets. We propose a hierarchical approach to clustering and classification that can accommodate very large datasets and tissues composed of complex mixtures of cell types and subtypes. For classification, our framework relies on support-vector machines (SVMs) with a linear kernel, trained on PCA-transformed data. To accurately distinguish between closely related cell types, we leverage our previously developed *kNN-smoothing* algorithm^11^ to effectively reduce technical noise levels. We demonstrate the capabilities of our framework by constructing and validating classifiers for human peripheral blood mononuclear cells (PBMCs), which allow the accurate prediction of immune cell types and subtypes in data generated using different scRNA-Seq protocols, as well as following experimental perturbation. We also apply our framework for construct a classifier for human pancreas cell types, demonstrating its general applicability.

## Results

### A scalable framework for robust scRNA-Seq cell type classification

To design a scalable framework for constructing robust cell type classifiers from large, heterogeneous scRNA-Seq datasets, we observed that cell types in a given tissue are often closely related biologically. For example, all subtypes of mature T cells found in peripheral blood originate from thymocytes, and can be expected to share similar transcriptomes. We therefore reasoned that a hierarchical approach would be most appropriate for cell type classification. In the example, a classifier could first generally distinguish T cells from the remaining cell types, and then further distinguish among the T cell subtypes. Importantly, this would allow the two classifiers to operate at different levels of resolution, and to utilize distinct sets of genes for classification, which would be more difficult to accomplish using a single classifier. With this hierarchical approach in mind, we designed effective and integrated methods for clustering, classification, and validation, which together comprise the *Moana* framework (**Figure 1a**).

**Figure 1:**
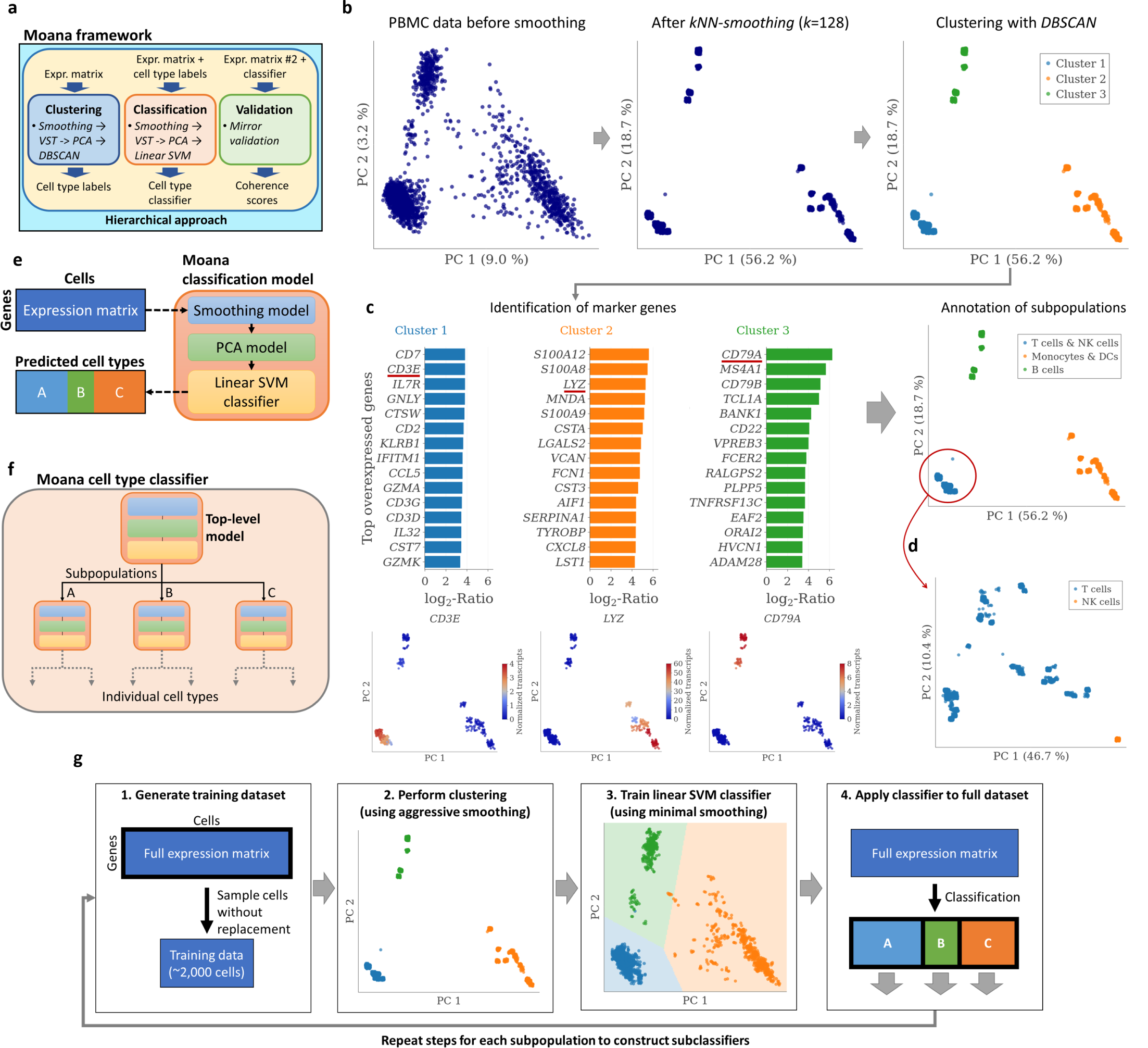
Overview of the Moana framework. **a** Schematic of framework components with input and output data (VST=variance-stabilizing transformation). **b** Clustering example using 2,000 cells from the PBMC-8k dataset (see text for details). **c** Annotation of cell types proceeds by identifying known markers among the most overexpressed genes for each cluster (top left). As an example, expression patterns of the *CD3D*, *LYZ*, and *CD79A* genes are shown (bottom left), which are known to be highly expressed in T cells, monocytes, and B cells, respectively. **d** Identification of additional subpopulations is accomplished by repeating the clustering procedure on cells from each cluster. **e** Schematic of the internal structure of the Moana classification model. **f** Schematic of the internal structure of a Moana cell type classifier, consisting of a hierarchy of independent classification models. **g** Strategy for efficiently performing clustering and classification for large datasets. A classifier is trained on a random subset of approx. 2,000 cells, and used to predict the remaining cells in the dataset. The procedure is repeated recursively for each subpopulation, enabling the construction of a complete hierarchical classifier without ever having to perform clustering or SVM training on the entire dataset.

Training of a machine learning classifier requires a dataset in which expression profiles are associated with cell type labels. As such labels are typically unavailable, it is necessary to first perform clustering with the goal of labeling each cell with a cell type or subpopulation. Given the high levels of technical noise, clustering of scRNA-Seq data can present a difficult challenge, for which numerous methods have been proposed^13^. For the purposes of our framework, we designed a new clustering method (**Figure 1b**) that takes advantage of our hierarchical approach by partitioning the cells into a few clearly distinct subpopulations, instead of resolving all cell types simultaneously. First, we aggressively smooth the data using our previously developed *kNN-smoothing* algorithm^11^, thereby removing as much technical noise as possible. We then perform clustering directly in two-dimensional PC space using *DBSCAN*, a standard clustering algorithm^18^. This allows a direct visual inspection of clustering results, and an examination of the expression patterns of known cell type markers (**Figure 1c**). Subsequently, additional subtypes can be identified by repeating the procedure for each subpopulation (**Figure 1d**).

Our clustering results suggested that with sufficient smoothing, even closely related cell types would become linearly separable in principal component space. However, overly aggressive smoothing can also introduce artifacts. In our framework, we therefore predict cell types using linear support vector machines (SVM), trained on minimally smoothed and PCA-transformed data (**Figure 1e**). Each principal component captures an expression module consisting of many genes, thereby reducing the impact of any single gene on the classification outcome. Additionally, SVM rely on the principle of maximum margin classification to achieve optimal classification performance^19^. Our classification model is therefore designed to exhibit maximal robustness with respect to protocol-related differences or unforeseen perturbations to the transcriptome. The “chaining” of individual classification models results in a hierarchical classifier that is able to accommodate a large number of cell types and subtypes in a given tissue (**Figure 1f**).

Datasets consisting of more than 10,000 cells present computational challenges related to the time and memory required to complete each of the analysis steps. We found that our hierarchical approach to constructing cell type classifiers allowed for a straightforward solution to this problem (**Figure 1g**). First, we generate a training dataset by randomly sampling a small subset of cells from the entire dataset. Clustering and training of a Moana classification model is performed on this training dataset, and the resulting classifier is then used to predict the cell types in the entire dataset. The procedure can then be repeated for each subpopulation, which allows the construction of a complete hierarchical classifier without ever having to analyze the full dataset. A detailed technical description of all components of the Moana framework is provided in the Methods.

### Construction and initial validation of a cell type classifier for human PBMCs

We applied our framework to construct a cell type classifier for human peripheral blood mononuclear cells (PBMCs), which present a difficult classification challenge owing to the relatively low RNA content of these cells (**Supplementary Figure 1**), and the fact that several cell types such as CD14+ and CD16+ monocytes are closely related, and are therefore expected to share similar transcriptomes. Using a PBMC dataset provided by 10x Genomics (PBMC-8k, *n*=8,381 cells), we constructed a Moana classifier comprising eight binary classification models, thus distinguishing between nine immune cell types: CD14+ monocytes, CD16+ monocytes, dendritic cells (DCs), B cells, NK cells, and four subtypes of T cells (**Figure 2a** and Methods). Our clustering method successfully identified subpopulations in the clustering steps required for the construction of this classifier (**Supplementary Figures 2 and 3**), demonstrating its effectiveness at different levels of resolution. While broadly distinguishing between T/NK cells, B cells, and Monocytes/DCs did not require any smoothing (*k*=1), smoothing was essential to accurately distinguish between most cell types in the training data (**Figure 2a** and **Supplementary Figure 4a**).

**Figure 2:**
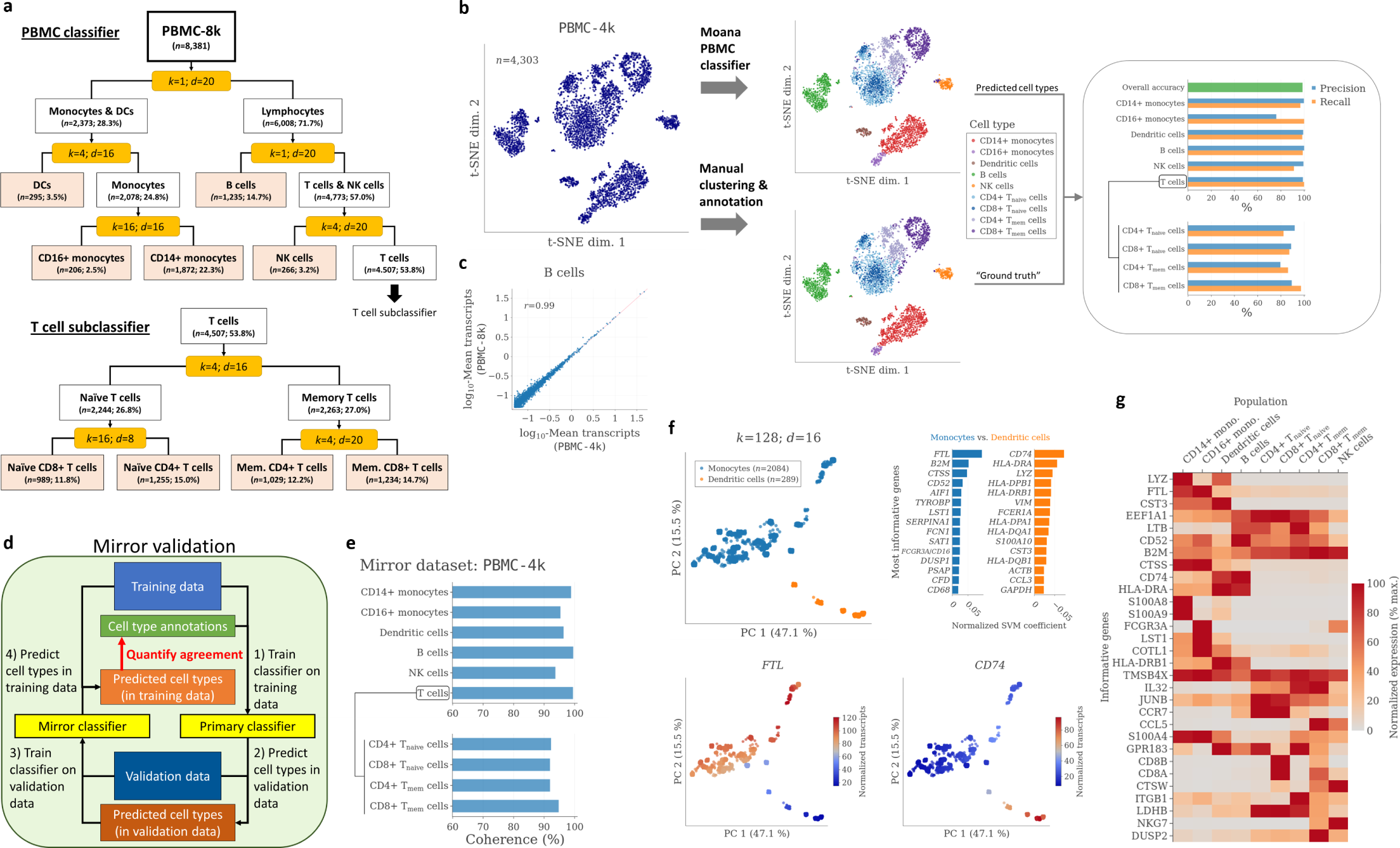
Construction and initial validation of a Moana cell type classifier for human PBMCs. **a** Classification tree of the classifier, trained on the PBMC-8k dataset. Yellow boxes represent classification models and indicate training parameters, *k* represents the number of neighbors used for smoothing, and *d* represents the number of principal components used. All models were trained using *ν*=0.02. The classifier distinguishes between nine cell types (shaded boxes). **b** Left: t-SNE analysis of all cells from the PBMC-4k dataset. Center: Cell type predictions (top) and annotations (bottom), overlaid onto the same t-SNE analysis. Right: Prediction accuracies, calculated by treating the annotated cell types as the ground truth. **c** Comparison of B cell average expression profiles in the two datasets. **d** Schematic overview of the mirror validation procedure. For large datasets, the cell types annotations (green box)can be replaced by the cell type identities predicted by the primary classifier. **e** Mirror validation results. **f** Clustering results (top left) and identification of most informative genes (top right) for the Monocyte/Dendritic cell classification model. Bottom: Smoothed expression levels of the *FTL* and *CD74* genes. **g** Heatmap showing the expression of the most informative genes from each classification model across cell types.

To validate our classifier, we obtained a second 10x Genomics dataset (PBMC-4k, *n*=4,340; cells obtained from the same donor as for the training dataset). We clustered the cells into the same nine cell types as before (**Supplementary Figure 5**), and manually excluded 37 cells that we identified as plasmacytoid dendritic cells (**Supplementary Figure 4c**). We then applied our PBMC classifier to this dataset, and used the clustering results as the “ground truth” for assessing the accuracy of the cell type predictions (**Figure 2b**). Prediction accuracies were generally high, with precision scores of above 98% for CD14+ monocytes, dendritic cells, B cells, and NK cells. Approx. 20% of cells predicted to be CD16+ monocytes were annotated as CD14+ monocytes during clustering. CD14+ and CD16+ monocytes did not form distinct clusters in t-SNE space, suggesting that the difficulty in accurately distinguished between them could be explained by their high transcriptomic similarity. While precision scores for T cell subtypes were slightly lower, ranging between 80-90%, this was still a remarkable result given that the naïve CD4+ and CD8+ T cell populations were completely overlapping in t-SNE space (**Figure 2b**). We used the predicted cell types to calculate average expression profiles for each cell type, which revealed near-perfect correlations between the training and the validation data (**Figure 2c** and **Supplementary Figure 4d**). These results demonstrated the ability of our classifier to accurately distinguish between closely related immune cell types in a validation dataset.

Our approach of combining PCA with linear SVM classification allowed us to quantify how informative each gene was for distinguishing between individual subpopulations (**Figure 2f** and Methods), which fundamentally depends on its cell type specificity as well as its absolute expression level. We collected the most informative genes from all subclassifiers and visualized them as a heatmap (**Figure 2g** and **Supplementary Figure 6**), which showed that these included both widely expressed genes such as *B2M* (Beta-2-microglobulin), as well as genes with highly cell type-specific expression such as *S100A8* and *S100A9* (expressed only in CD14+ monocytes and dendritic cells). This demonstrated that by choosing a hierarchical approach to cell type classification, we were able to utilize information from many more genes than if we had tried to construct a classifier based solely on genes with highly cell type-specific expression patterns.

### Systematic assessment of classification performance in the absence of a ground truth

The generation of a “ground truth” through manual clustering can be time-consuming and carries the risk of introducing subjective biases or mistakes, which can result in the over- or underestimation of classification accuracies. Unfortunately, a more reliable ground truth, such as scRNA-Seq data from experimentally purified subpopulations, is frequently unavailable. To overcome this limitation, we devised an algorithm which we refer to as *mirror validation* (**Figure 2c** and Methods). In this approach, we use the trained classifier to predict cell types in a validation dataset from the same tissue, and then use the predicted cell type labels to train a new “mirror classifier” on this dataset. We then apply the mirror classifier to the original training dataset, and assess the degree with which the predictions from mirror classifier agree with those of the original classifier. We reasoned that whenever the original classifier fails to accurately identify its cell types in the validation dataset, it would not be possible for the mirror classifier to accurately recapitulate the original cell type assignments in the training dataset. This lack of “coherence” would therefore indicate a failure to accurately determine cell types in the validation dataset.

We applied mirror validation using PBMC-4k as the validation dataset, and found that coherence scores, which combine precision and recall values into a single measure (Methods), were at or above 90% for all cell types (**Figure 2e**), suggesting high classification accuracies. By examining the validation results for individual classification models, we observed that the validation accuracies for highly imbalanced classes (e.g., T cells and NK cells) could be further improved to over 98% by lowering the *ν* parameter of the mirror classifier (**Supplementary Figure 4b**). However, independent of the choice of *ν*, the accuracies for the classifiers distinguishing monocyte and T cell subtypes remained slightly lower than for the other cell types. Therefore, the mirror validation results closely agreed with the results obtained using the manually generated “ground truth”, demonstrating that mirror validation allowed the effective assessment of classification performance in the absence of a ground truth.

### Accurate prediction of PBMC cell types in data generated using a different scRNA-Seq protocol

Given the diversity of scRNA-Seq protocols developed^4^, scRNA-Seq classifiers should ideally be able to accurately determine cell types in data generated using a different protocol, which may result in substantial systematic expression differences. We therefore aimed to assess the ability of our classifier to accurately predict PBMC cell types for data generated using 10x Genomics’ “v1” chemistry, as opposed to the “v2” chemistry used to generate PBMC-8k. A t-SNE analysis of a combined dataset with both v1 and v2 PBMC expression profiles showed that even after normalization, cells clustered by dataset, not cell type, suggesting the presence of large systematic differences that we did not observe between datasets generated using the same protocol (**Figure 3a** and **Supplementary Figure 7d**). When we applied our classifier to the v1 PBMC dataset (v1-PBMC-16k, *n*=16,000), cells of all cell types were identified, although cell type proportions differed significantly from the v2 training data (**Figure 3b** and **Supplementary Figure 7a**). Using the predicted cell types, we calculated average expression profiles for each cell type, and found that the protocol-dependent differences indeed appeared larger than many cell type differences (**Figure 3b** and **Supplementary Figure 7b, c**).

**Figure 3:**
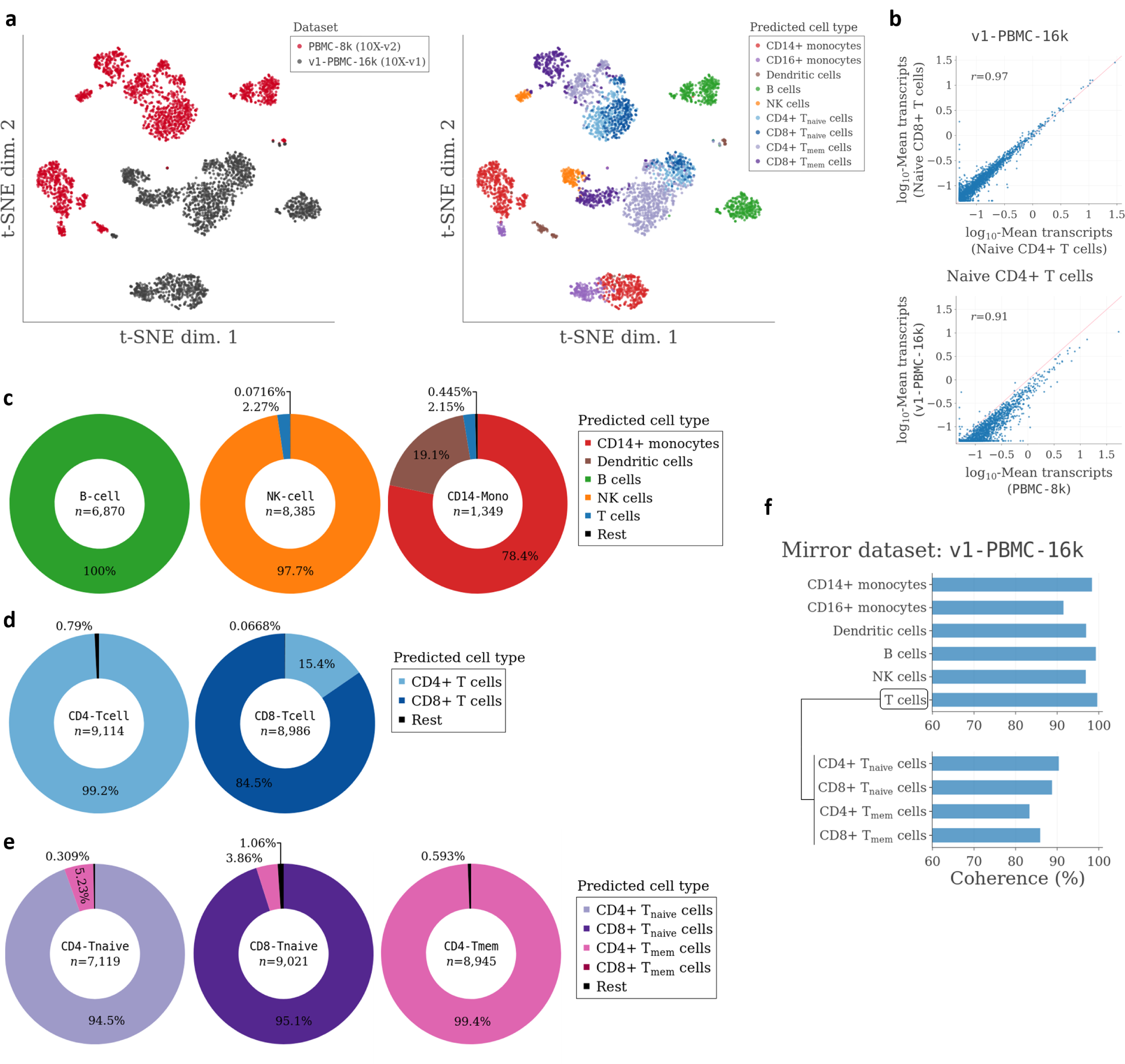
Accuracy of the Moana PBMC classifier for data generated using a different scRNA-Seq protocol. **a** Left: t-SNE analysis of 4,000 cells, consisting of a 50:50 mixture of cells from PBMC datasets generated using two different protocols (10x Genomics v1 vs. v2 chemistry). Right: Same t-SNE analysis, overlaid with the cell type identities predicted using the PBMC classifier (**Figure 2a**). **b** Top: Comparison of average naïve CD4 and naïve CD8 T cell expression profiles in the v1 dataset. Bottom: Comparison of average naïve CD4+ T cell expression profiles in the v1 and v2 datasets. **c-e** Prediction results of the PBMC classifier, for v1 datasets representing experimentally purified PBMC subpopulations (Methods). The “Rest” category encompasses any cell type not listed in the legend, as well as cell types with a predicted abundance of less than one percent. **f** Mirror validation results for the PBMC classifier, using v1-PBMC-16k as the validation dataset.

The availability of v1 datasets of purified PBMC subpopulations^20^ allowed us to use an experimentally generated ground truth to assess the accuracy of our classifier. The classifier identified B cells, T cells and NK cells with >97% accuracy, but predicted the presence of 19% dendritic cells in a dataset representing purified CD14+ monocytes (**Figure 3c**). We did not explore the extent to which this could reflect an experimental contamination associated with the negative selection protocol used by Zheng et al.^20^, as opposed to a classification error. We next applied the classifier to data from experimentally purified T cell subpopulations, and found that more than 99% of total CD8+ T cells were predicted to belong to the CD8+ lineage, while 85% of total CD4+ T cells were predicted to belong to the CD4+ lineage (**Figure 3d**). In addition, we found that classification accuracies ranged from 94-99% for naïve CD4+, naïve CD8+, and memory CD4+ T cells (**Figure 3e**). Overall, these results demonstrated the ability of our classifier to accurately distinguish between closely related cell types in data exhibiting substantial systematic differences associated with the use of a different scRNA-Seq protocol.

### Moana outperforms a previously described method for scRNA-Seq cell type prediction

To directly compare our classification results to those obtained with a previously proposed classification method, we reimplemented the *scmap-cluster* method^17^, a non-hierarchical approach that relies on gene selection and correlation measures to predict cell type identities. We first ensured that we were able to accurately reproduce the classification results from the original study (**Supplementary Figure 8**). We then trained an *scmap-cluster* classifier on the PBMC-8k dataset, while using the PBMC-4k dataset to determine the optimal number of genes (Methods). We then applied this classifier to the experimentally purified PBMC subpopulations. We found that B and NK cells were predicted accurately, while accuracies were much lower for all other cell types (**Supplementary Figure 9**). In particular, for all purified T cell subpopulations, the classifier left between 28% and 70% of cells “unassigned”, indicating an inability to conclusively determine cell type identities. These results showed that the *scmap-cluster* classifier failed to achieve a similar resolution as our Moana classifier when presented with data that exhibited substantial batch effects.

### Construction and validation of a classifier for human pancreas cells

Moana is a general framework for constructing cell type classifiers for complex tissues based on scRNASeq data. To demonstrate its applicability beyond PBMC samples, we constructed a human pancreas cell type classifier based on scRNA-Seq data reported by our lab (Baron16- 3, *n*=4,591; Methods).^21^ We visually confirmed the presence of batch effects, as well as the ability of our classifier to predict cell types independent of those effects (**Figure 4a**). Cell type composition was highly variable across cell types (**Figure 4b**), but mirror validation showed that cell type predictions were highly coherent, thereby validating the accuracies of our predictions (**Figure 4c**). Previous work relied on well-established marker genes to identify individual cell types, and we found that our predictions allowed us to recover those marker genes in an unbiased fashion (**Figure 4d** and **Supplementary Figure 10a**), further confirming our classification results. Finally, since some of these marker genes exhibited very high expression levels, we asked if it would be possible to predict cell type identities based on only a single marker gene for each cell type. We found that this was the case for the highly specialized endocrine cell types, but not for stellate and ductal cells (**Figure 4e**). In contrast, among PBMCs, we were only able to predict B cells using a single marker gene (**Supplementary Figure 10b,c**). Our results confirmed the general applicability of our framework, and showed that while distinguishing between the endocrine cell types was indeed a trivial task, our classifier also enabled the accurate prediction of the remaining cell types in the pancreas.

**Figure 4:**
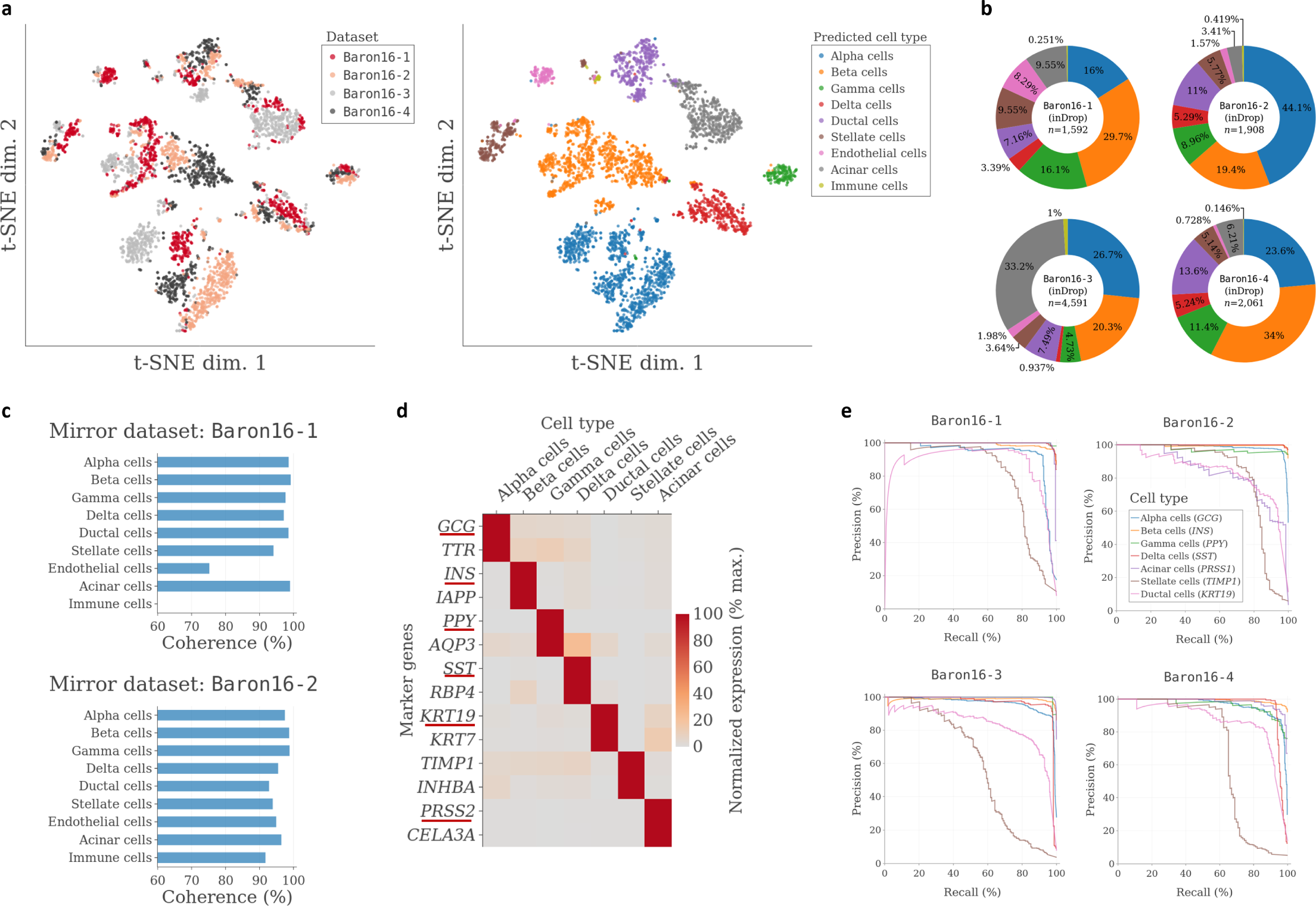
Prediction of human pancreas cell types. **a** Left: t-SNE analysis of 4,000 cells consisting of an equal mixture of cells from four independent biological samples^21^. Right: Same t-SNE analysis, overlaid with the cell type identifies as predicted from a *Moana* classifier trained on Baron16- 3. **b** Predicted cell type proportions for each of the four samples. **c** Mirror validation results, using two different samples as mirror datasets. **d** Heatmap showing the expression of cell type marker genes identified based on the predicted cell type identities. Genes that were used as cell type markers by Baron et al.^21^ are underlined. **e** Precision-recall curves for identifying individual cell types based on the expression of a single marker gene for each cell type (as indicated in the legend).

### Accurate prediction of PBMC cell types after an experimental perturbation

In addition to exhibiting robustness to batch effects, an scRNA-Seq cell type classifier should also be able to identify cell types following in vivo or in vitro perturbations to the “normal” cell state. To test whether a Moana classifier would be able to accurately identify cell types following an experimental perturbation, we decided to predict cell type identities for PBMCs following exposure to the cytokine interferon beta (IFN-β)^22^. This treatment was reported to result in widespread transcriptomic changes across cell types and the downregulation of key cell type markers^15,23^, thus providing an interesting test case for the robustness of Moana classifiers. We first used mirror validation to test the performance of our original PBMC classifier on the unexposed control dataset (Kang18-Ctl, *n*=12,757), which indicated that it did not accurately distinguish between CD14+ and CD16+ monocytes, nor between T cell subtypes (**Supplementary Figure 11a**). We observed that the unexposed cells exhibited dramatic transcriptomic differences relative to our original training data (**Supplementary Figure 11c**), potentially resulting from their culturing for the duration of the experiment^22^. Evidently, these differences were too great to allow an accurate classification of subtypes using our original PBMC classifier. We therefore decided to use the control dataset to train a new PBMC classifier (**Supplementary Figure 12** and Methods). The structure of the classifier largely mirrored that of our first PBMC classifier, however we were unable to accurately distinguish T cell subtypes using our clustering method, perhaps owing to the significantly lower transcript counts in this dataset (**Supplementary Figure 1a**). We then applied this classifier to predict the cell types in the treatment dataset (Kang18-Tx, *n*=13,551).

We visually confirmed the presence of significant expression differences between treated and untreated cells (**Figure 5a**), and found that our classifier identified cells from all major cell types with similar proportions in both conditions, in agreement with previous analyses of the same data^15,22^ (**Figure 5a,b**). We next used the treatment dataset to perform mirror validation, which resulted in high coherence scores for all cell types except for dendritic cells (**Figure 5c and Supplementary Figure 11b**). We also compared our cell type predictions to the cell type assignments from a previous analysis by Butler et al., who performed clustering after “alignment” of the two datasets^15^, and found that for both datasets, our predictions strongly agreed with these annotations, with the exception of dendritic cell predictions in the treatment dataset (**Supplementary Figure 11d**). As dendritic cells represented only 2% of cells in the training data, it is possible that additional parameter tuning was necessary to achieve better classification robustness for these cells. Overall, these results demonstrated the ability of a Moana classifier to accurately predict cell types following an experimental perturbation that induced widespread transcriptomic differences.

**Figure 5:**
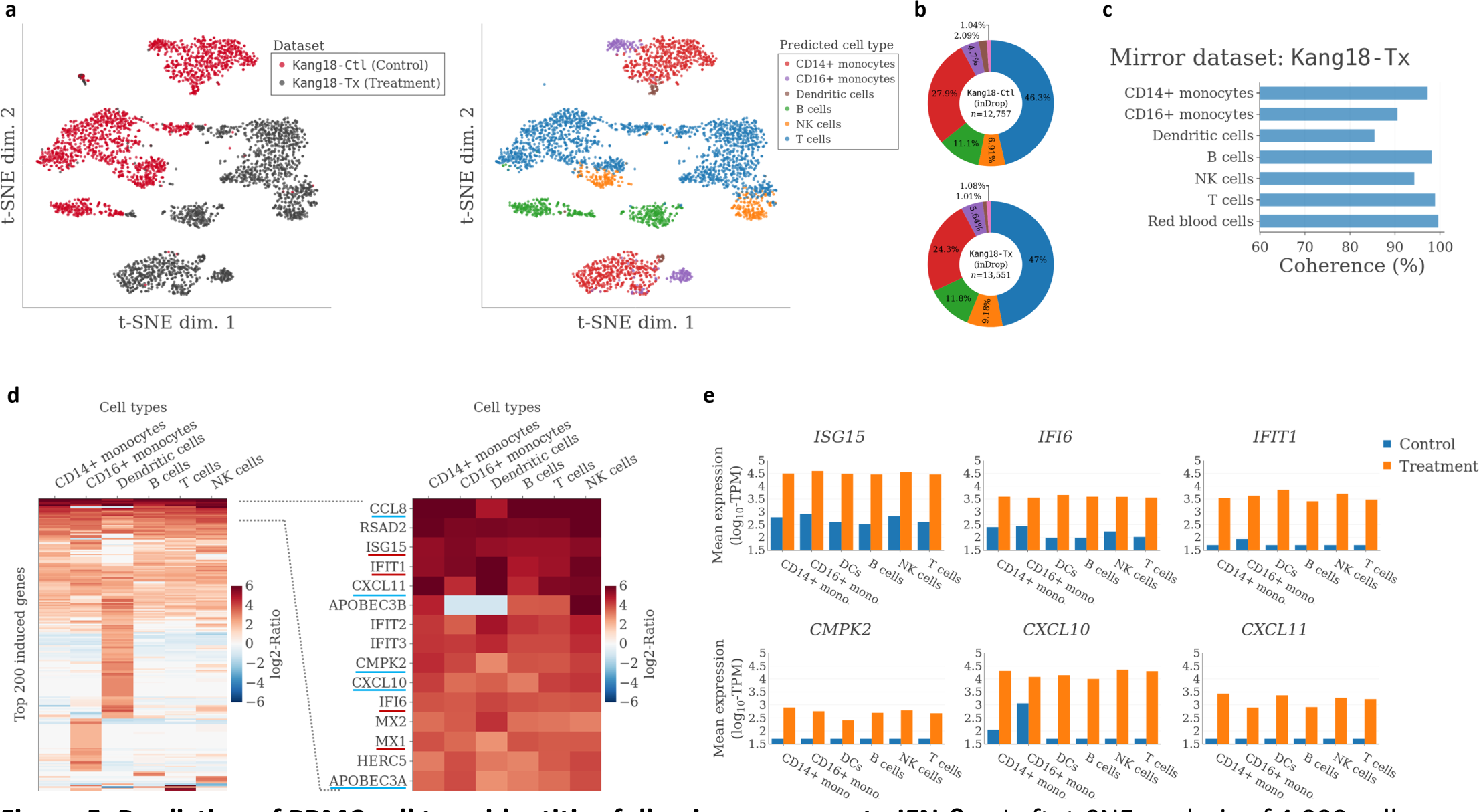
Prediction of PBMC cell type identities following exposure to IFN-β. **a** Left: t-SNE analysis of 4,000 cells consisting of an equal mixture of cells from the control and treatment datasets, respectively^22^. Right: Same t-SNE analysis, with predicted cell types overlaid. **b** Predicted cell type proportions. **c** Mirror validation results. **d** Left: Heatmap showing cell type-specific gene expression levels after IFN-β exposure, relative to their maximum expression level across cell types in the control condition. Right: Zoomed-in view of 15 genes with very strong upregulation. Underlined are genes reported as either universal (red) or cell type-specific (blue) IFN-β response markers by Butler *et al.*^15^. **e** Detailed view of expression levels with and without IFN-β exposure for three “universal” (top) and three “cell type-specific” (bottom) marker genes.

### Expression analysis based on predicted PBMC cell types suggests a broadly shared response to IFN-β

Finally, we aimed to use the predicted cell type identities as the basis for a cell type-specific analysis of differential expression following exposure to IFN-β. A previous analysis that applied clustering to “aligned” datasets had reported a number of genes that were strongly induced across all cell types, as well as other genes with more cell type-specific induction^15^. We defined a “physiological” expression level for each gene as the maximum expression level across all cell types in the control, and then compared the expression levels in the treatment condition to this level. Our results suggested the presence of at least two distinct responses to IFN-β (**Figure 5d**): First, a shared response that involved a strong upregulation of more than 50 genes in all cell types, and second, a similarly strong dendritic cell-specific response. Since dendritic cells (DCs) represented a rare population and we did not distinguish between plasmacytoid and myeloid DCs, we decided to focus our analysis on the shared response. Unexpectedly, we found that in addition to genes previously reported as universal IFN-β markers, this response encompassed several genes such *CXCL11*, *CCL8* and *CMPK2* that were previously reported as having a cell type-specific response^15^ (**Figure 5d, e**). As the analysis was performed on the same dataset, using closely matching cell type assignments (**Supplementary Figure 11d**), this discrepancy likely resulted from different approaches to quantifying differential expression. Overall, these results demonstrated that Moana cell type predictions allowed the systematic quantification of cell type-specific differential expression following exposure to IFN-β, and suggested that independent of cell type, PBMCs elicit a common response that results in the strong upregulation of a substantial number of genes.

## Discussion

In this work we have introduced *Moana*, a generally applicable machine learning framework that enables the construction and deployment of accurate and robust cell type classifiers for high-throughput scRNA-Seq data. We applied this framework to construct PBMC classifiers, which we were able to show allowed the accurate prediction of cell types in data generated using different protocols, as well as following perturbations of the transcriptome. To our knowledge, this is the first demonstration of an accurate and robust scRNA-Seq cell type classifier for PBMCs, and we propose that future benchmarks of scRNA-Seq classification performance should focus on difficult challenges presented by PBMCs and other tissues that contain small- to moderately-sized cells and closely related cell types. During the preparation of this manuscript, Alquicira-Hernández et al.^24^ proposed a classification framework that shares some basic building blocks with Moana, in that it involves the training of SVM classifiers on PCA-transformed data. Despite substantial differences in methodology, we think that the independent proposal of combining PCA with SVM for scRNA-Seq cell type classification speaks to the intuitive appeal of this approach. However, the scope of the current work significantly exceeds that of previous reports17,24. We describe a hierarchical framework that encompasses methods for clustering, classification and validation, all of which directly integrate a specialized smoothing algorithm to reduce technical noise levels in scRNA-Seq data^11^. Moreover, we show that our framework can be used to construct classifiers that exhibit high prediction accuracies for a tissue composed of multiple closely related cell types, even in the presence of strong batch effects.

Our hierarchical approach to clustering and classification represents a key innovation, as it simultaneously enables our framework to scale to large datasets, accommodate a large number of cell types, and perform clustering and classification at distinct levels of resolution. Another key advantage of this approach is the ability to construct individual subclassifiers using different datasets. For example, a subclassifier for T cell subtypes could be trained on a different scRNA-Seq dataset containing only T cells. By collecting scRNA-Seq data after enriching for specific subpopulations, it is thus possible to experimentally overcome the computational difficulty of training (sub-)classifiers for extremely rare populations of cells. In this work, we have proposed to remedy the often widely differing cell type abundances by creating synthetic training datasets in which rare cell types are overrepresented, which helps to increase classification robustness as long as a certain minimum number of cells of a particular type are available.

In addition to extremely rare subpopulations, we decided to ignore the presence of doublets (multiplets) in the data. The ability of PCA to capture specific sources of variation in individual PCs allowed us to construct accurate classifiers without removing doublets. However, their identification is important to avoid confusion during the clustering stage, and we would ultimately like to incorporate an automated method for removing these technical artifacts from the data in our framework. We briefly considered modeling doublets as artificial cell types, however this approach did not seem appropriate given the fact that the transcriptomic composition of the doublets depends entirely on the actual cell type composition of the sample. We would instead favor a simulation-based approach such as the one proposed by McGinnis et al.^25^ Additionally, to further increase the ability to accurately distinguish between closely related cell types or states, future research may be directed at the selection of informative principal components^24^, and at the incorporation of a gene selection step^26^ during both clustering and classification, as PCA does not admit a sparse data representation.

In advancing quantitative cell type definitions, we are not advocating for a rigid or mindless application of those definitions. Rather, cell type classification can form the basis for an exploratory analysis of scRNA-Seq data, including a search for entirely new cell types. Moana classifiers are designed to provide maximum robustness to batch or treatment effects. As the nature and magnitude of those effects are unknown in advance, it would be very difficult to maintain the desired robustness while simultaneously trying to identify cells that represent a previously unseen cell type. However, cell type classification results can be followed up with clustering or outlier detection analyses aimed at identifying such new cell types. A distance threshold for suggesting the presence of a new cell type would perhaps be best defined empirically, based on an analysis of many datasets with the same classifier, and the observed distribution of similarities with the cells in the training data.

Our work was motivated by the large number of completed or ongoing “atlas-building” projects^21,28–33^, in which large scRNA-Seq datasets are being produced for individual organisms or tissues, with the goal of generating comprehensive cell type compendia. Moana provides a way to distill these datasets into robust cell type classifiers that can be applied to automatically determine cell types in new scRNA-Seq datasets. New and powerful single-cell RNA-Seq protocols are producing datasets comprising billions of noisy gene expression measurements^1^. It is increasingly clear that it will not be possible to effectively extract and organize biological knowledge from these measurements without equally powerful computational frameworks and tools. In designing Moana, we aimed to develop a framework that was tailored to the noise characteristics of scRNA-Seq data, relied as much as possible on established machine learning methods (e.g., PCA, SVM), allowed the direct visual examination of intermediate results, and maintained computational efficiency. Given the numerous biological and computational challenges involved, we believe that the collaboration of experimental and computational biologists is essential in realizing the full potential of scRNA-Seq technology.

## Methods

A description of all methods is contained in the Supplement.

## Acknowledgements

We would like to thank Yun Yan for processing the pancreas raw data with the SingleCell pipeline. In addition, we would like to thank Dr. Gal Avital, Dr. Gustavo S. França, Dr. Eva Chmielnicki, Bo Xia, Felicia Kuperwaser, Reuben Moncada, and Dalia Barkley for their comments on an earlier version of this manuscript.

